# Membrane binding promotes oligomerization and functional activation of viral RNA-dependent RNA polymerase

**DOI:** 10.1101/2025.09.08.674988

**Authors:** Oshri Afanzar, Nanqin Mei, Ingrid Altunin, Priscilla L. Yang

## Abstract

Positive-sense, single-stranded RNA (ssRNA^+^) viruses initiate infection by utilizing host ribosomes to directly translate proteins from their genomes. One of these proteins is the RNA-dependent RNA polymerase (RdRp), which is responsible for replicating the viral genome. Hepatitis C virus (HCV), a well-studied ssRNA^+^ virus, encodes a membrane-bound RdRp, here termed NS5B, that can form oligomers. NS5B exists in two conformations; only one is believed to be capable of binding RNA. The relationship between membrane localization, oligomerization, and conformation remains unclear. We investigated whether membrane localization mediates NS5B’s oligomerization, conformation and function. In solution, NS5B exhibited temperature-dependent oligomerization that correlated with cooperative RNA binding and a conformational shift at micromolar concentrations. In contrast, total internal reflection fluorescence microscopy and atomic force microscopy revealed that membrane-bound NS5B forms stable, non-terminating oligomers at nanomolar concentrations, often localized near lipid raft domains. The presence of oligomers was correlated with membrane recruitment of RNA and nucleotides, suggesting functional activation. The lower concentration threshold for RNA binding on membranes may be explained by membrane-induced oligomerization that promotes the RNA-binding conformation of NS5B. Our findings are consistent with a model in which membrane association regulates NS5B’s function through oligomerization-driven conformational control.

## Introduction

The *Riboviria* realm of viruses is characterized by the presence of an RNA-dependent RNA polymerase (RdRp) encoded in their genomes and includes many human pathogens(1). Diseases caused by these viruses are amongst the leading causes of death in the United States(2), and their economic impact on livestock and poultry further underscores their significance. Within this realm, positive-sense single-stranded RNA (ssRNA^+^) viruses directly translate their RNA genomes into proteins using host ribosomes. This includes the RdRp and accessory proteins responsible for synthesis of a complementary negative-strand RNA that in turn serves as a template for producing new viral genomes (see Chapter 5 in reference for an overview of viral replication strategies(3)). Experimental models suggest that the switch between transcription and translation is regulated(4–6)(Figure 1A), but the underlying mechanism(s) regulating for transcription initiation by viral RdRps is an open question.

**Figure 1.**
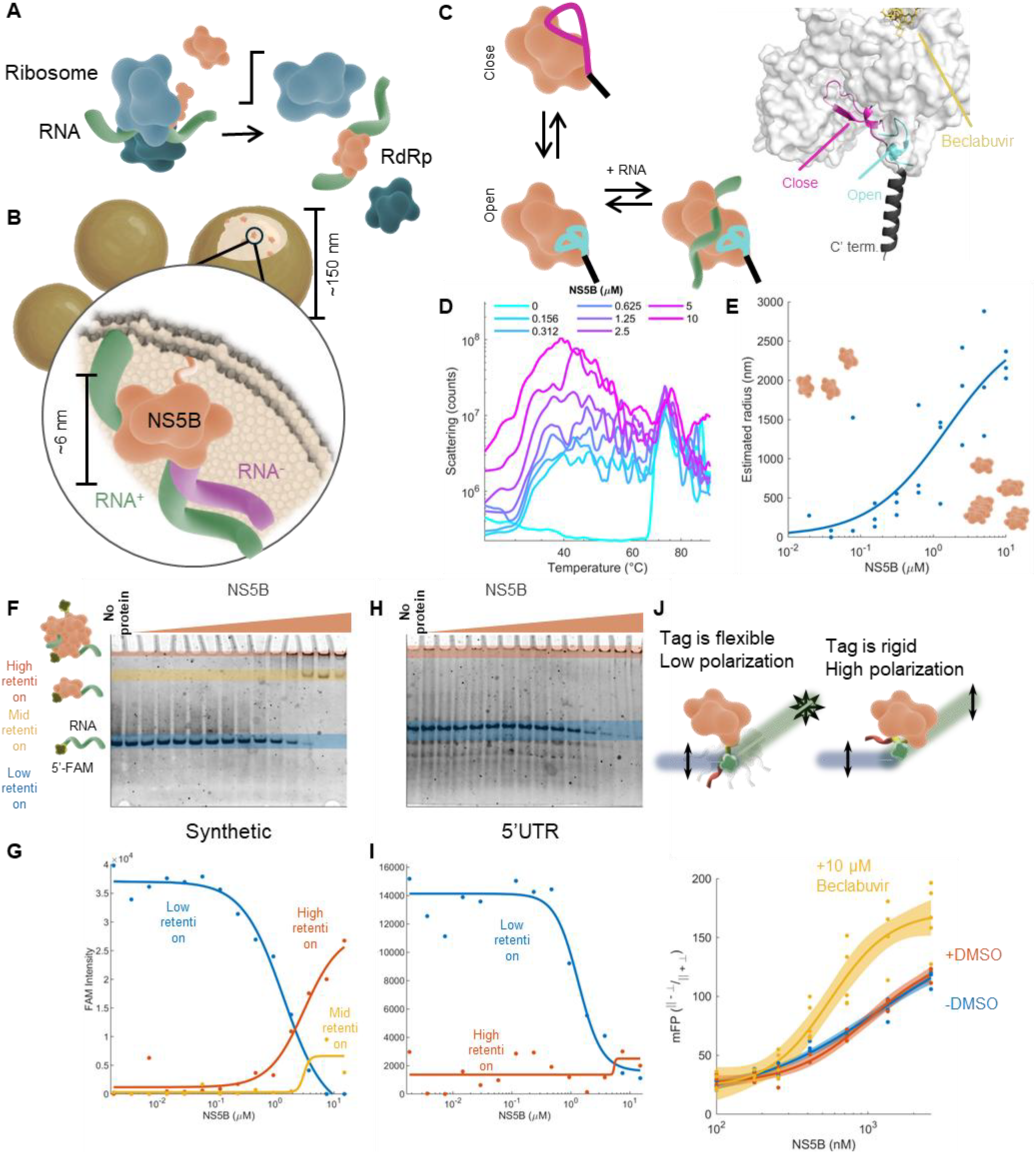
NS5B, the membrane-tethered RdRp of HCV, functionally oligomerizes in a concentration and temperature-dependent manner. **A.** Positive-sense single-stranded RNA (ssRNA^+^) viruses like HCV deliver their genomes into the host cytoplasm, where translation begins immediately. At later stages, the translated viral RdRp initiates RNA replication, marking a “transcriptional switch.” **B**. In HCV, RNA replication is mediated by membrane-tethered RdRps (NS5B) localized to virus-induced membranous compartments. **C**. NS5B exists in “closed” and “open” conformations with potential functional relevance. *Left*: A schematic model where RNA binding (green) requires a conformational switch of NS5B. *Right*: NS5B AlphaFold structure demonstrates the predominantly closed NS5B conformation where upstream to the C-terminal membrane tether (black), C^reg^ (magenta) folds toward the protein core. C^reg^ conformation fold was validated against PDB: 1YUY. In the presence of the allosteric NS5B inhibitor becalbuvir (yellow), C^reg^ adopts an exposed conformation (cyan; PDB:4NLD) . **D**. Light scattering suggests NS5B oligomerization in concentration- and temperature-dependent manner. The experiment was replicated 4 times with similar outcomes. Representative data are shown. **E**. Dynamic light scattering measurement of NS5B at 37 ºC. Each data point is an independent experiment. The line is a fitted Hill model with K_0.5_ ∼ 1.5 µM. **F-I**. Native PAGE shows that RNA binding is correlated with oligomerization. Synthetic (F) or 5′UTR (H), 5′-FAM-labeled RNA at 75 nM was incubated with increasing NS5B concentrations at 37 °C for 30 min and resolved on gel. Shaded areas indicate RNA migration: unbound RNA (blue) and RNA complexes (red and yellow). Fluorescence intensity at the shaded regions was quantified for the Synthetic (G) or 5′UTR (I) experiments. Data represents >3 independent experiments. **J**. Fluorescence polarization shows NS5B’s C-terminal correlates with oligomerization. *Top:* when the FlAsH-labeled C-terminus is mobile, polarization is low; restricted motion yields higher polarization. *Bottom:* Polarization of 100 nM FlAsH-labeled NS5B upon titration with increasing unlabeled NS5B. Each point represents the average of 2-4 technical replicates from a single experiment. Lines represent Hill fits; shaded areas denote 95% confidence intervals, fitted for K_D_ ± 95% CI: 1.1 ± 0.5 µM (buffer), 1.0 ± 0.3 µM (DMSO), and 0.56 ± 0.14 µM (becalbuvir).

The *Flaviviridae* Family is composed of ssRNA^+^ viruses associated with various human and vertebrate diseases(7). Hepatitis C virus (HCV), a significant member of this family, causes chronic liver disease and has been a major contributor to virus-related morbidity worldwide (8). Despite the availability of curative antiviral therapies(9, 10), some patient groups show poor responses to treatment or access to treatment. Consequently, the disease remains prevalent(11), with approximately 10,000 chronic HCV-related deaths and 100,000 new chronic patients reported annually in the U.S.(12) and similar rates globally(13). The search for HCV antivirals has generated valuable biological insights and experimental tools, providing a foundation for understanding viral mechanisms and potentially developing future therapeutic strategies for ssRNA^+^ viruses.

Cell membranes play a crucial role in HCV replication. Like many other ssRNA^+^ viruses(14), HCV forms specialized membrane compartments that house its membrane-tethered viral RdRp, nonstructural protein 5B (NS5B), to facilitate genome replication (Figure 1B)(15–17). NS5B is tethered to membranes through a hydrophobic C-terminal domain consisting of the last 21 amino acids of the protein(18). While membrane association is not necessary for NS5B’s *in vitro* polymerase activity, it is essential for viral replication, suggesting that membrane interactions regulate its function(18, 19). Regulation by membrane binding has been implicated in several key biological pathways(20–22), indicating it may be a fundamental principle. Although the role of NS5B’s membrane tethering has not been extensively studied, recent findings suggest that NS5B activity increases when reconstituted on a supported lipid bilayer(23). We have previously shown that HCV-infected cells accumulate desmosterol, a cholesterol precursor, due to cleavage of 24-dehydrocholesterol reductase (DHCR24) by the HCV NS3-4A protease, and that this change affects HCV RNA replication(24–26), supporting the idea that membrane lipid composition is also important for RNA replication. Desmosterol enhances membrane fluidity in synthetic models and HCV replication membranes (27), leading to the hypothesis that it may improve the mobility of viral proteins in cellulo and in vivo. Importantly, desmosterol accumulation does not reduce cholesterol levels, which are also vital for replication(24, 28).

The structure of NS5B resembles a right-hand shape typical of RdRps with palm, fingers, and thumb domains and exists in two conformations(29). In the predominant “closed” conformation(30, 31), association of double-stranded RNA (dsRNA) seems unplausible due to narrowness of the RNA-binding cavity, and rearrangement to the “open” conformation is thought to allow RNA binding(29, 32). However, NS5B, like other polymerases(33), is likely to bind RNA in an “open” state and switch back to a “closed” state while maintaining the RNA bound for processive elongation. One of the determinants of the “open” conformation is rearrangement of the 21-50 residues upstream of the C-terminus, here termed “C^reg^”, but also known as the “β-flap” (34–37). C^reg^ is thought to interfere with dsRNA binding (38) and its role is apparent from mutations and deletions in this region that enhance NS5B’s activity (19, 39, 40). C^reg^-mediated regulation is further supported by studies showing that beclabuvir, a compound that induces an “open” conformation, shifts C^reg^ from a buried to solvent-exposed state(32) (Figure 1C). Furthermore, truncation of C^reg^ was shown to affect inhibitor-mediated protein stability (41). A mechanism to explain how these changes in NS5B’s conformation are regulated is not yet known.

RdRp oligomerization is consistently observed in many ssRNA^+^ viruses (42–49). HCV’s NS5B oligomerization has been reported, but a functional role for oligomerization has not been established(50–54). One possible outcome of oligomerization is cooperativity which can be broadly defined as the emergence of function as a result of allosteric homo-interactions (recently reviewed herein(55)). Thus, oligomerization may promote NS5B cooperative function as conformation switching. Here, we investigated how NS5B’s oligomerization affects its function and whether membrane localization regulates these processes.

## Results

We hypothesized that the processes of oligomerization and membrane binding are linked to the conformation and autoregulation of NS5B. HCV has six genotypes, which differ in their genetic sequences but share similar mechanisms of replication(56). For our studies we utilized a recombinant NS5B protein derived from genotype 2a in which the hydrophobic C-terminal 21-residue membrane tether was replaced with FlAsH and 6×His tags to enable site-specific fluorescent labeling, affinity purification, and affinity-based membrane tethering. We refer to this construct as NS5B(Δ21)-FlAsH-His.

### NS5B liquid phase oligomerization correlated with RNA binding and conformation at micromolar concentrations

To assess the oligomerization state of purified NS5B(Δ21)-FlAsH-His, we first analyzed its molecular weight distribution using size-exclusion chromatography. At an estimated concentration of 1 µM, the protein eluted predominantly as a monomer at 4 °C (SI Figure 1). Although NS5B is known to oligomerize, optimal conditions for this process have not been defined. We therefore hypothesized that oligomerization may occur more readily at higher concentrations and physiologically relevant temperatures. To test this, we monitored light scattering by the protein, a property that increases with the formation of high molecular weight species across a range of concentrations and temperatures. Because protein denaturation at elevated temperatures reduces light scattering, a temperature-dependent decrease helps distinguish oligomers from aggregates that may form during non-physiological denaturation. We found that all protein concentrations showed a sigmoidal increase in light scattering between 30-40 °C, consistent with accelerated oligomerization under physiological conditions. This was followed by a decrease above 40 °C, consistent with loss of oligomers due to denaturation. This interpretation is supported by a parallel increase in fluorescence ratio above 40 °C measured by differential scanning fluorimetry, independently indicating protein denaturation (SI Figure 2). To further assess oligomerization, we used dynamic light scattering at 37 °C to estimate average particle size across concentrations. This revealed apparent cooperative oligomerization of NS5B (K_D_ ∼ 0.75 µM, n ∼ 3 from a Hill model fit). Together, these results support the interpretation that NS5B forms oligomers at micromolar concentrations and physiological temperatures in the liquid phase. Accordingly, all subsequent experiments were conducted at 37 °C to support oligomer formation.

After we observed NS5B concentration-dependent oligomerization we hypothesized that oligomerization is associated with protein function. One possibility is that oligomerization may drive the switch to the open conformation to enable RNA-binding. To investigate this idea, we examined how NS5B’s RNA-binding changes in response to NS5B concentration. For this purpose, we used two different synthetic 5′-FAM-labeled RNAs: a 53-base synthetic hairpin based on a SARS-CoV-2 study model(57) (referred to as “synthetic”) and an 88-base fragment corresponding to the 5′ negative-strand of HCV’s 2a replicon(58) (referred to as “5′UTR”), both at a concentration of 75 nM. Samples were incubated with increasing concentrations of NS5B and then resolved on a native gel. Gel migration of the FAM-labeled RNAs did not change with increasing protein concentrations up to 300 nM (Figure 1F-I). Lack of gradual RNA binding can indicate that the protein was predominantly in a “closed” state. Above 300 nM, we observed binding of both RNAs by NS5B that was nonlinear, a switch consistent with oligomerization-mediated cooperativity. In the case of the synthetic hairpin RNA, the nonlinear shift in RNA migration manifested in three distinct forms. First, increasing NS5B concentration resulted in a nonlinear depletion of free RNA (Figure 1F-G, blue), which correlated with the formation of a slower-migrating RNA species at three-digit nanomolar to micromolar protein concentrations (Figure 1F-G, red). At 1-10 µM concentrations, an additional faster-migrating RNA signal appeared, suggesting the formation of lighter molecular weight protein-RNA complexes (Figure 1F-G, yellow). The non-linear transition in RNA migration pattern is consistent with cooperative behavior and may result from oligomerization of NS5B. For the 5′UTR, which is the natural substrate for NS5B, we observed a nonlinear reduction in free RNA at supramicromolar NS5B concentrations. High molecular weight bands, including the yellow bands in Figure 1F were generally absent for this model. Because this model is double in size compared to the synthetic model, this is possibly due to large protein-RNA complexes failing to enter the gel (Figure 1H-I). The NS5B concentration range that altered RNA migration in the gel was aligned with concentrations previously observed to affect RNA migration in size-exclusion chromatography and with RNA fluorescence polarization measurements at increasing NS5B concentrations(54, 59). For both RNAs, RNA binding occurred at a similar concentration to that required for protein oligomerization in the absence of RNA. This argues that oligomerization itself might facilitate RNA binding by inducing the closed-to-open transition of the polymerase

To obtain additional evidence for oligomerization-dependent conformational shifts, we employed an established fluorescence polarization approach (60) to assess the local conformation of NS5B’s C-terminus at varying protein concentrations. It should be emphasized that conformation change that is associated with increasing protein concentration is a cooperative function, which necessitates inter-protein interaction. We measured the fluorescence polarization of NS5B-tagged with a FlAsH fluorophore at the C-terminus. At a concentration of 100 nM, the polarization values indicated a mobile C-terminus (Figure 1J, blue; this should not be confused with a mobile C^reg^ which is upstream to the label). To determine if NS5B oligomerization affects protein conformation, we added unlabeled NS5B to the FlAsH-labeled protein. We observed an increase in fluorescence polarization as the concentration of unlabeled protein increased (Figure 1J, blue). These results are consistent with the idea that NS5B oligomerization promotes a conformational shift in the C-terminus, although they could also be explained by oligomerization stabilizing or limiting the mobility of the C-terminus or FlAsH tag. To confirm that increased protein concentration promotes an “open” conformation, we measured the effect of beclabuvir, a compound known to induce an “open” state conformation in NS5B and C^reg^(32). As expected, we observed that beclabuvir shifted the response curve for RNA-binding to lower NS5B concentrations. No effect was noted at 100 nM protein concentrations (Figure 1J, yellow), which is consistent with the possibility that NS5B oligomerization is necessary for beclabuvir’s action. This can also indicate that beclabuvir binds and stabilizes NS5B in its “open” conformation. Overall, our findings show that NS5B’s RNA-binding is associated with a conformational switch which is protein concentration dependent.

### Tens to hundreds of nanomolar NS5B makes stable, non-terminating oligomers on the face of membrane

Building on previous evidence that membrane-tethering enhances NS5B activity(23), we investigated whether it also promotes oligomerization. We created a supported lipid bilayer system containing desmosterol and incorporating Ni-NTA lipids to allow tethering of His-tagged NS5B. FlAsH-labeled NS5B was introduced at increasing concentrations (0-5 µM) in a flow chamber, allowing unbound protein to be washed away. Control experiments using only FlAsH dye showed a weak and easily removable signal, confirming minimal nonspecific binding. In contrast, FlAsH-labeled NS5B produced distinct, immobile fluorescence spots (Figure 2A). The absence of lateral protein diffusion on membranes in which lipids were mobile indicated the formation of oligomers, as membrane mobility is expected to decrease with increasing molecular weight. These spots first appeared at approximately 50 nM NS5B and increased sharply in number and intensity at around 500 nM (Figure 2B-C). This transition likely reflects higher-order assembly or aggregation. Since NS5B’s membrane-binding is limited by the availability of Ni-NTA in these experiments, NS5B concentrations above 500 nM must involve interactions between membrane-bound and free NS5B (see SI Methods), suggesting that oligomerization extends beyond direct membrane tethering.

**Figure 2.**
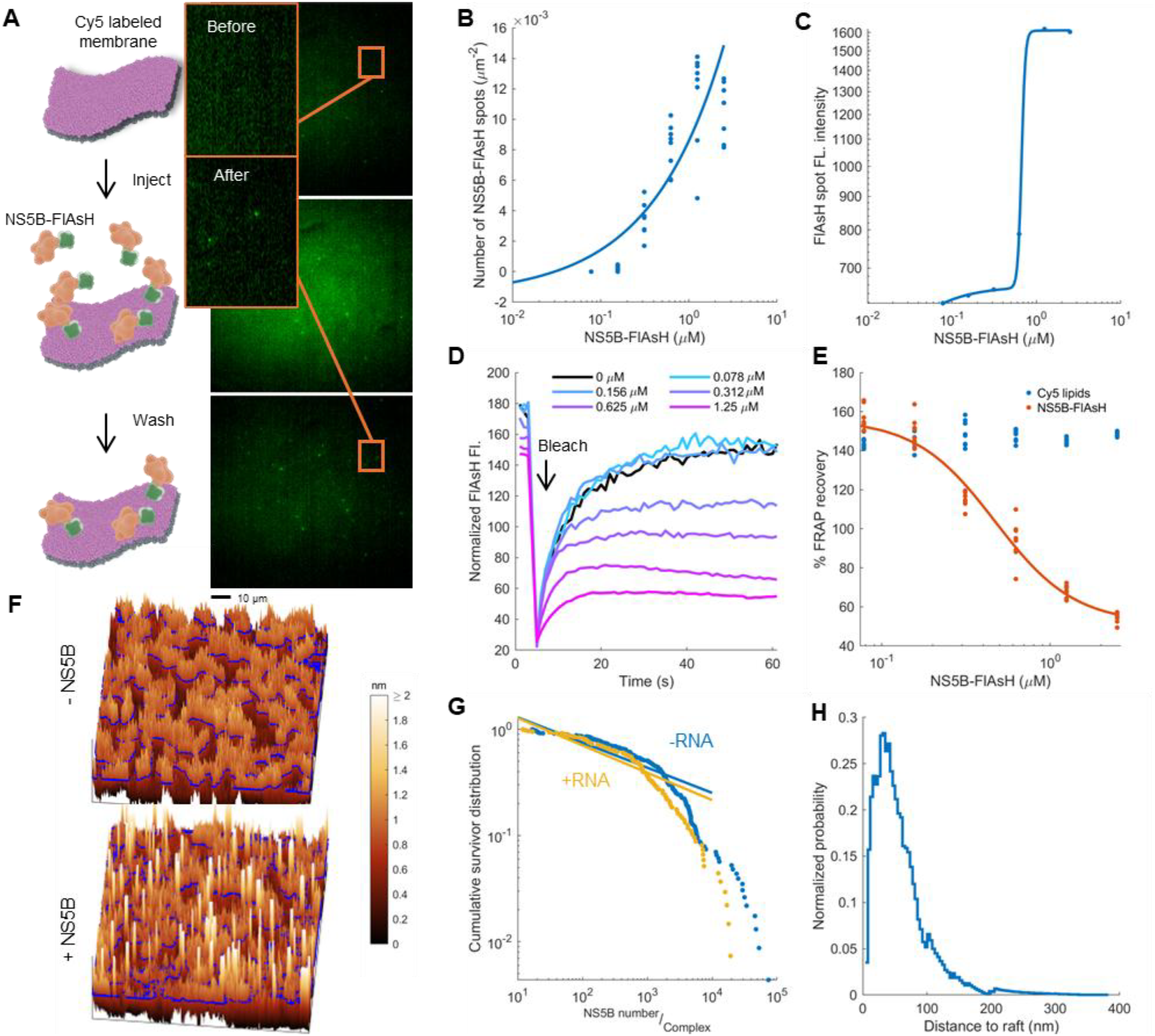
NS5B makes stable, non-terminating oligomers on the face of membrane. **A.** NS5B forms membrane immobile spots. *Left*: Experimental schematics where FlAsH-labeled NS5B is introduced to supported lipid bilayer containing Ni-NTA and Cy5 head groups modified lipids, to enable immobilization of NS5B(Δ21)-FlAsH-His and monitor membrane integrity, respectively. *Right*: Representative FlAsH fluorescence images before, during, and after injection of ∼50 nM FlAsH labeled NS5B. **B**. Quantification of all visible membrane-associated NS5B spots at increasing FlAsH-NS5B concentrations. Each point represents quantification of all spots from a single field of view from the same experiment. Three full experiments were performed (two in flow chamber, one in wells) and three partial ones (flow chamber; covering <500 nM concentrations). **C**. Average fluorescence intensity per spot from the same dataset as in B. Line shows a bi-Hill fit as a visual guide. with additional partial replicates. **D**. NS5B membrane mobility assessed by FRAP. *C*urves show normalized FRAP of FlAsH-labeled NS5B that was applied in increasing concentrations to the same membrane. Each trace is an average of eight acquisitions. **E**. Endpoint FRAP signal quantification from D. Data shown for FlAsH-labeled NS5B (red) and Cy5-labeled membranes (blue), from the same samples used in B-D. **F**. Representative atomic force microscopy scan of 3 µm^2^ membrane in the absence (top) or presence (bottom) of 200 nM NS5B. Domains that were < 2nm elevated from lipids, possibly representing sterol domains, outlined in blue; height is color-coded (see color bar). **G**. Cumulative survival distribution for NS5B domains volume from a 9 µm^2^ scan. Line is fitted cumulative power-law distribution. Data is representative. Similar results were observed in three independent experiments. **H**. Cumulative distances between NS5B- and sterol-domains for experiment as in G, adjusted by subtracting sterol-to-non-sterol distance distributions as a density control.

To differentiate between dynamic and stable membrane-associated oligomers, we employed fluorescence recovery after photobleaching (FRAP) on FlAsH-labeled NS5B. A high-intensity 405 nm laser was used to bleach specific membrane regions, and recovery was monitored over one minute. Fluorescent lipid controls showed full recovery across all conditions. In the absence of NS5B, fluorescence was minimal and fully recovered, likely due to signal bleed-through. At concentrations below 150 nM NS5B, fluorescence intensity increased and recovery was complete, indicating high membrane mobility (Figure 2D-E). This is consistent with an interpretation that much of the NS5B population exists in diffusive, low-molecular weight forms. However, at higher concentrations, recovery progressively declined, suggesting that a growing fraction of NS5B became immobile and tightly bound as oligomers or aggregates. Some minor recovery persisted even at the highest concentrations, indicating a residual mobile (likely monomeric) population. The lack of trend toward full fluorescence recovery at higher concentrations implies the formation of stable oligomers or aggregates. Assuming a diffusion coefficient of 0.1 µm^2^/s, we estimated the dissociation constant between monomeric and oligomeric/aggregated NS5B to be less than 10 nM (see SI Methods). These results suggest that NS5B forms tightly bound, low-dissociation oligomers/aggregates on membranes in a concentration-dependent manner.

NS5B membrane formations may exhibit two types of oligomerization: terminating oligomerization, where the arrangement of subunits limits growth (similar to capsid assembly), and non-terminating oligomerization, such as fibril growth which lacks a defined endpoint. The former is expected to produce oligomers of distinct sizes, while the latter may not have a defined size. To differentiate between these two scenarios, we examined changes in membrane topography using atomic force microscopy (AFM) in the presence and absence of NS5B. Supported lipid bilayers were prepared on mica substrates, and we imaged areas of approximately 3 µm^2^ at an x-y resolution of about 6 nm in both the presence and absence of 100-200 nM NS5B. In the absence of the protein, the membrane exhibited height variations typical of sterol and rigid lipid phase separation, with sterol raft-like domains elevated by less than 2 nm (Figure 2F, up). After introducing NS5B, distinct topographic features approximately 6 nm in height and varying in lateral dimensions emerged (Figure 2F, down). Assuming NS5B monomers volume of 216 nm^3^ (taken as minimal x-y dimensions of AFM resolved objects and assuming height of 6 nm; NS5B’s volume is ∼110 nm^3^) we calculated that these features represent NS5B oligomers following a power law-like distribution in the number of monomers, ranging from tens to tens of thousands of monomers per oligomer (Figure 2G; solid lines represent a power law fit to the empirical cumulative distribution data). This power law-like distribution suggests that a significant portion of the protein mass is concentrated in a small fraction of high molecular weight formations. These formations are likely not aggregates because they would be expected to show disordered growth in all dimensions. As expected for oligomers, we observed the height of the membrane-associated NS5B domains was within 4-7 nm, suggesting monolayer oligomers of lateral association. The addition of RNA, whether synthetic or 5’UTR, did not produce a distinguishable or consistent effect. This lack of effect is consistent with the interpretation that RNA-bound NS5B detaches from the oligomer to form a mobile complex that is unlikely to be detected by AFM (see Discussion). Notably, oligomers tended to localize near the boundaries of sterol rafts more frequently than expected by chance (Figure 2H). However, it remains unresolved whether this proximity is due to selective recruitment to phase-separated Ni-NTA lipids or reflects an intrinsic raft-targeting behavior of NS5B.

### Membrane-bound NS5B binds RNA and nucleotides at nanomolar concentrations

Since NS5B’s RNA-binding activity is correlated with oligomerization and protein conformation in the liquid phase (Figure 1), we next examined whether RNA-binding follows the appearance of NS5B oligomers in protein concentrations that cannot support RNA-binding in liquid phase. To determine this, we monitored NS5B’s concentration-dependent RNA-binding on a supported lipid bilayer formed in PDMS wells. NS5B was labeled with ReAsH, a red-shifted analog of FlAsH(60), allowing for spectral separation from the FAM-labeled RNA (either synthetic or 5’UTR) used to estimate RNA binding. We applied NS5B at concentrations ranging from approximately 1 to 100 nM, washed the samples, and imaged them in both ReAsH and FAM fluorescence channels before and after the addition of RNA (Figure 3A).

**Figure 3.**
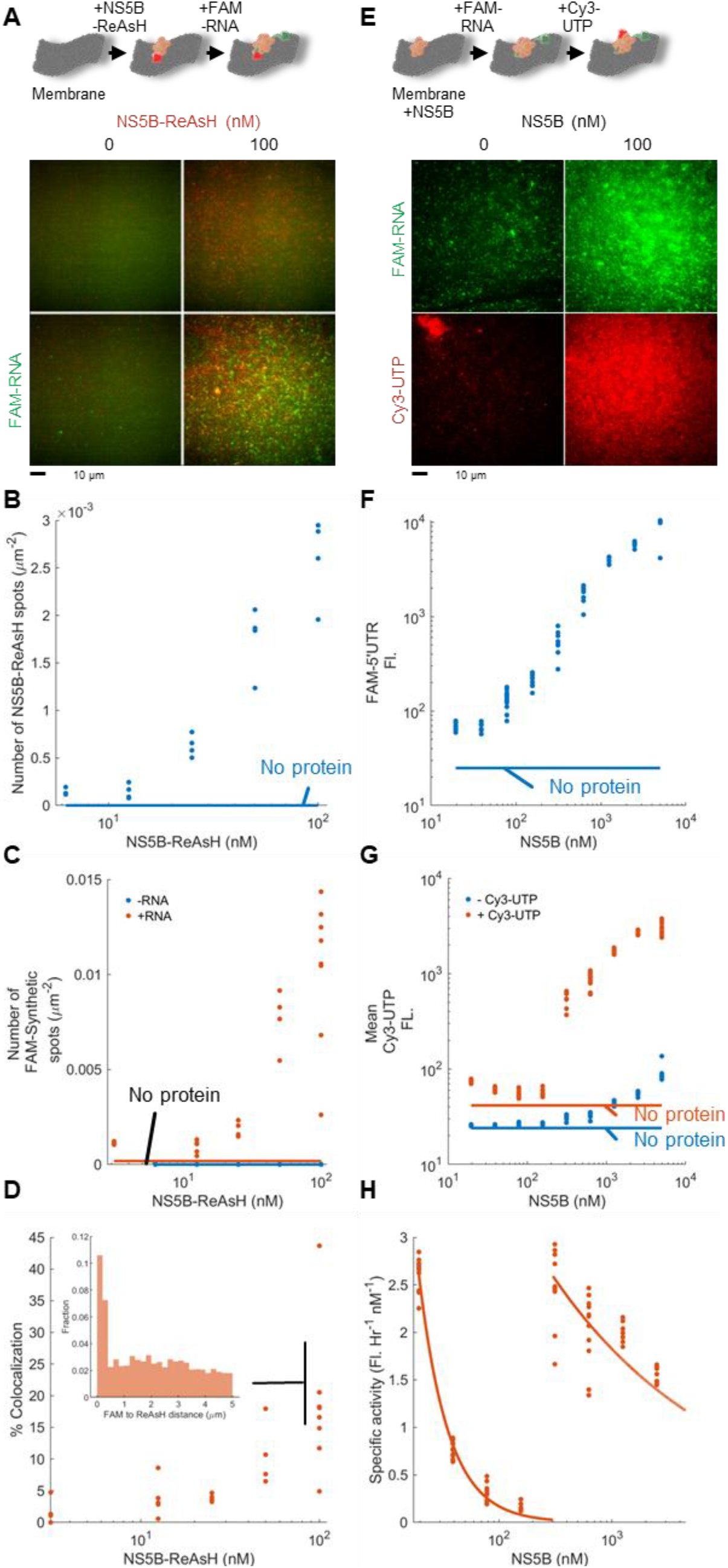
Membrane-tethered NS5B oligomers recruit RNA and nucleotides at nanomolar concentrations. **A-C**. Detectable formation of RNA binding oligomers of NS5B initiates at nanomolar concentrations. Experimental setup description (A, top) and representative fluorescence montage (A, bottom) showing membrane tethered NS5B(Δ21)-ReAsH-His in the presence or absence of 100 nM FAM-labeled synthetic RNA. Signals for NS5B (B) and RNA (C) were quantified before (blue) and after (red) the addition of RNA. Horizontal lines indicate mean signal in the absence of NS5B. Data is representative of three experiments, in one of which 5′UTR-was employed instead of synthetic-RNA. **D**. The fraction of FAM-to-ReAsH colocalization (≤500 nm distance), was determined based on data in B-C. Inset: FAM-to-ReAsH distance distribution at 100 nM NS5B. **E-G**. Recruitment of ribonucleotides to the membrane is NS5B-dependent. Experimental setup description (E, top) and representative fluorescence montage (E, bottom) showing membrane-tethered NS5B (100 nM) in the presence or absence of 5′UTR FAM-RNA (1000 nM) and/or Cy3-UTP/NTPs (200 μM Cy3-UTP, 1 mM each NTP). Quantification of membrane-associated RNA (F) and UTP (G) was determined as a function of NS5B concentration, before (blue) and after (red) Cy3-UTP incubation. Horizontal lines indicate the average of a no-NS5B control. Data is representative of three experiments. Some RNA signals was bleeding into the Cy3 channel before the addition of Cy3 (blue data in panel G; note the log scale). **H**. Data from G normalized to NS5B concentration to approximate catalytic efficiency. Red lines are visual guides. Similar results were obtained in three independent experiments.

The number of ReAsH-NS5B fluorescence spots increased in a concentration-dependent manner becoming pronounced at the initial concentration of 10 nM NS5B and higher (Figure 3B). The subsequent addition of FAM-labeled RNA resulted in a corresponding increase in the number of membrane-associated FAM spots. FAM-labeled RNA was not observed when NS5B was omitted, indicating that the presence of NS5B results in RNA recruitment to the membrane (Figure 3C). Spatial analysis of the fluorescent spots revealed that only about 15% of the overall population showed colocalization between FAM and ReAsH signals (Figure 3D). This observation, along with the NS5B-mediated import of RNA to the membrane and the lack of perfect colocalization with NS5B oligomers is consistent with the interpretation that some RNA-NS5B oligomers are too small to show as distinct spots on the membrane.

To assess whether membrane-bound NS5B can show a catalytically active conformation after RNA loading, we tested its ability to recruit nucleotides to the membrane surface. Unlabeled NS5B was allowed to bind to supported lipid bilayers in PDMS wells and was subsequently incubated with FAM-labeled 5’UTR RNA. After washing out unbound components, we introduced a nucleotide mix containing 20% Cy3-labeled UTP and incubated the samples for 1 hour at 37 ºC before removing excess reactants. The experiment included NS5B concentrations ranging from nanomolar to micromolar (Figure 3E shows an example with 100 nM NS5B). As expected, RNA recruitment increased proportionally with NS5B concentration, indicating that the association of RNA with membranes is dependent on NS5B (Figure 3F; the horizontal blue line represents the no NS5B control). In all experiments involving Cy3-UTP, we observed that the Cy3 signal localized to the membrane in an NS5B-dependent manner.

In contrast to RNA recruitment, Cy3-UTP recruitment exhibited a biphasic pattern at NS5B concentrations below 100 nM. Between 10 and 100 nM NS5B, the Cy3-UTP signal on the membrane increased relative to the no NS5B control (Figure 3G, horizontal red line for no NS5B control), but paradoxically decreased with increasing protein concentration, suggesting a regime of negative cooperativity (Figure 3G, red data points for measurements with NS5B). At higher concentrations (hundreds of nM to low µM), Cy3-UTP membrane association increased again, this time in a nonlinear manner. Notably, no UTP-Cy3 incorporation was observed in the RNA polymerization assays conducted in solution and analyzed by urea-PAGE assays (Data not shown). Assuming the Cy3 signal observed with membrane-associated NS5B reflects non-saturating transcriptional activity, we estimated NS5B’s catalytic efficiency across these concentration regimes. To do this, we plotted the Cy3-UTP signal divided by NS5B concentration, providing an approximation of catalytic turnover efficiency. This analysis revealed reduced catalytic efficiency at both low and high NS5B concentrations (Figure 3H). Similar results were obtained when the data were normalized to RNA signal to account for non-saturating RNA binding by NS5B (Data not shown). Although, it is not clear why two regimes exist or what their physiological relevance is, though they may relate to membrane formed oligomers at up to hundreds of nanomolar and liquid formed oligomers/aggregates at increased concentrations. The apparent diminished shift in catalysis for each regime may indicate a relationship between oligomerization levels and the probability of NS5B performing a successive transcription initiation cycle. Additional investigation is required to distinguish these possibilities.

## Discussion

In this study, we investigated how RNA-binding, ribonucleotide binding, and possibly RNA synthesis are influenced by NS5B oligomerization and membrane-binding. We observed that oligomerization is correlated with the NS5B functions of RNA binding and C-terminal conformation in solution at micromolar concentrations (Figure 1). We found that NS5B forms stable oligomers with non-terminating topology on membrane surfaces nearby sterol rafts (Figure 2), starting at concentrations where oligomerization was hard to distinct in solution (compare Figure 3B with Figure 1E). Consistently, membrane-localized NS5B could bind RNA at an order of magnitude lower concentration than was required for binding in solution (compare Figure 3C, F with Figure 1F-I). We also observed a nonlinear relationship between Cy3-UTP localization to the membrane surface and NS5B concentration in the presence of RNA (Figure 3G). This can indicate a switch between open and closed NS5B conformations to mediate both RNA-binding and RNA synthesis (Figure 3F-H); higher resolution measurements are needed to examine this explicitly. We acknowledge that our interpretation is limited by the varying detection sensitivities across different assays. Nonetheless, consistent with previous findings(23), we believe that membrane-localization regulates NS5B activity through oligomerization-mediated conformation switch.

To frame our findings in the perspective of membrane function in viral replication, we propose a model in which NS5B oligomerization acts as a concentration- and membrane-dependent molecular switch (Figure 4A). Newly translated NS5B is anchored to intracellular membranes, where its lateral diffusion is regulated by virus-induced changes in sterol metabolism. The accumulation of desmosterol may alter membrane properties or topology to favor oligomer nucleation. This lateral diffusion, which restricts NS5B orientation, should increase the frequency of productive collisions between monomeric NS5B, leading to higher oligomer formation rates than expected in solution. As local NS5B concentrations rise, transcriptionally competent oligomers form, particularly at lipid raft boundaries, which are believed to coincide with viral RNA replication sites. The transcriptional switch may be temporally gated to allow upstream processes such as NS3-4A-driven modulation of the cellular lipid environment and formation of replication compartments(15, 24, 25) to occur before robust RNA synthesis begins. Additionally, membrane association may prevent inappropriate transcription in the cytosol, thereby reducing the production of double-stranded RNA species that could trigger host immune responses.

**Figure 4.**
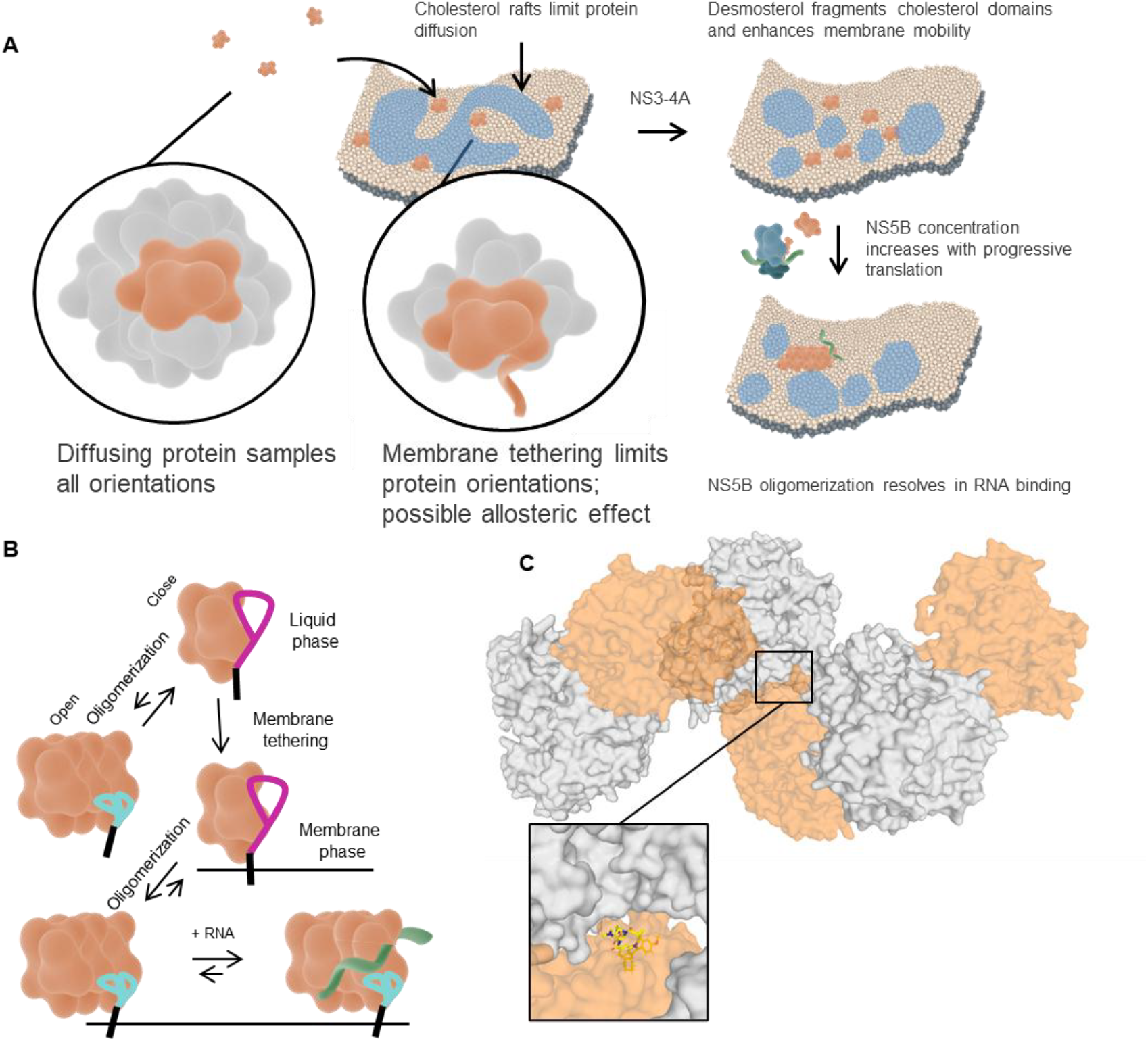
Hypothesis for the effect of membrane localization on transcription initiation in HCV. **A.** NS5B membrane localization by C-terminus tethering diffusion is hindered by cholesterol domains(27). Viral modulation of sterols enables NS5B diffusion, alongside continuous production of NS5B increases NS5B concentration and rate of self-association. This eventually lead to oligomerization and consequent binding of viral genomic RNA, activating transcriptional switch. **B**, Proposed role for NS5B oligomerization in regulating RNA binding. From left up to down right: NS5B exists in predominantly “closed” state monomer in liquid phase with poor oligomerization. Upon membrane-tethering, NS5B oligomerizes and switches to the “open” conformation that is competent to bind RNA. **C**, AlphaFold Multimer predicts non-terminating oligomerization of NS5B through two protein faces, one of which overlaps with beclabuvir binding site. Every other protein in the oligomer is shown as a semitransparent orange surface. The beclabuvir (yellow) binding interface is shown in the blow-out. Visualization by Pymol(63).

The molecular coordinates by which membrane association might affect NS5B conformation are not clear. We suggest that conformation shifts due to oligomerization regulate switching between closed and open states for the loading of RNA toward catalysis (Figure 4B). How NS5B oligomer initiates transcription from its RNA-bound state is not clear; single-molecule approaches will likely be required to resolve and investigate possible heterogeneity in subunit competence within the oligomer(61). To understand how oligomerization can lead to the “open” conformation, we generated an AlphaFold multimer prediction of NS5B. The resulting model resembles the crystal structure packing of poliovirus RdRp (Figure 4C). The predicted model indicates non-terminating oligomerization, with interfaces overlapping with the binding site for beclabuvir. Thus, we speculate that NS5B oligomerization stabilizes the “open” conformation by acting on the beclabuvir allosteric site. If confirmed, this mode of oligomerization may be a conserved mechanism of RdRp activity across ssRNA^+^ viruses.

While in vitro studies of isolated components, such as those described here, provide valuable mechanistic insights, they may not fully capture the spatial and temporal complexity of viral replication in cells, where multiple viral and host factors interact. Cell-based studies have been crucial for characterizing higher-level replication processes but are limited in resolving biochemical mechanisms. Future efforts to dissect viral transcription dynamics should prioritize the reconstitution of multi-component systems. One promising approach is the use of cytoplasmic sheet cell-free systems(62), which allow for high-resolution spatiotemporal observation of cellular dynamics in unconfined living cytoplasm over extended periods, making them well-suited for studying viral dynamics. Furthermore, the common modality of many RdRps to oligomerize on membranes surface raises the question of mechanistic conservation of the observations described herein.

## Materials and Methods

### Protein Production and Labeling

The NS5BΔ21-FlAsH-6×His construct, codon-optimized for E. coli (genotype 2a, synthesized by Twist Bioscience), was expressed in E. coli BL21 cells under kanamycin selection. Cells were induced at OD_600_ ∼0.6 with 0.2 mM IPTG for 6 h at room temperature, then lysed in “NS5B buffer” (400 mM NaCl, 50 mM NaPi pH 8.0, x1 ROCH protease inhibitor, ∼0.1 mg/ml DNase I). Lysates were clarified by centrifugation (40,000 rcf, 30 min), and protein purified by Talon (Co^2+^) affinity chromatography followed by heparin column using a 100-1000 mM NaCl gradient. Aliquots (20-60 μM) were stored at -80 °C in the presence of 10% glycerol. When indicated, protein was labeled with ≥2× molar excess FlAsH-EDT_2_ (MedChemExpress) or TC-ReAsH (Invitrogen); residual dye removed by PD-10 columns (Cytiva).

### Dynamic Light Scattering and nanoDSF

Protein samples diluted with NS5B buffer were loaded to high sensitivity DLS capillaries and measured by Prometheus Panta (NanoTemper).

### Fluorescence Polarization

FlAsH labeled protein at ≤100 nM was mixed with unlabeled protein diluted by NS5B buffer at different concentrations. For beclabuvir experiments, 10 μM drug was incubated with FlAsH-labeled NS5B for 2.5 h before mixing 1:1 with unlabeled NS5B. Fluorescence polarization was measured (BMG plate reader; ex: 482±16 nm, LP504 dichroic, em: 530±40 nm). Fluorescent probe concentration was set at twice the lowest concentration in linear fluorescence range; <10% of signal attributed to non-probe components.

### RNA Templates

5′ FAM-labeled RNA sequences were ordered from IDT:

5’UTR:

AUACUAACGCCAUGGCUAGGCGCUUUCUGCGUGAAGACAGUAGUUCCUCACAGGGGAGUGAUUCAUGGCGGAGUGUCGCCCCU

Synthetic: AGGGUUUUUUUUUUUAUUAUUUAUUCUUUAUUUUUCUUGCGUAGUUUUCUACG

### Native Gel RNA-Protein Binding

NS5B (0-15 μM) was incubated with constant RNA (75 nM) in 400 mM NaCl, 50 mM NaPi, 5 mM MgCl_2_, plus RNase inhibitor (1 μL/50 μL) at 37 °C for 30 min. Samples were mixed 1:1 with loading buffer and run on 10% native PAGE in 1× TBE (120 V, 40 min). Imaging: Cy2 channel on ImageQuant 800.

### Lipid Vesicle Preparation

Lipids (DPPC, DOPC, desmosterol, Cy5, DGS-NTA(Ni)) from Avanti Polar Lipids were mixed in chloroform at 3:3:2 (DPPC:DOPC:desmosterol), with optional 0.1% Cy5 and 4% NTA. After argon drying and vacuum desiccation, lipids were resuspended in buffer, sonicated (15 min), and extruded (100 nm filter). Vesicles (1 mg/mL) were stored at 4 °C and used within 5 days.

### Fluorescence Microscopy, Sample Preparation and Image Analysis

Microscopy was performed using a CellTIRF Olympus system (SI methods for full description). For sample preparation, coverslips (#1.5H, 24×50 mm) were cleaned with Piranha solution, plasma treated, and assembled with PDMS wells or flow chambers. Vesicles were deposited, incubated 30 min, and washed. Cy5-labeled membranes were FRAP-validated for mobility. For flow chamber experiments, PTFE-plumbed PDMS chambers were connected to computer-triggered syringe pump (NE-4000, 50-100 µL/min). NS5B (FlAsH-or ReAsH-labeled) was flowed in, incubated 2 min, then washed. For wells experiments, protein concentrations (total volume ∼50 µL) were incubated in PDMS wells coated with supported lipid bilayer at 37 °C for 30 min, washed (10-15×), and treated with RNA (100-1000 nM) and washed again. When indicated, membrane tethered protein-RNA complexes were subsequently incubated for 1 h in the presence of 200 µM Cy3-UTP and 1 mM ATP, GTP, and CTP. This was followed by additional washing. MATLAB (R2024b) was used for all analyses. For FRAP, Cy5 membrane channels were fit to hyperbolic recovery curves; samples of poor fits were excluded. Recovery curves were averaged after interpolation.

### Atomic Force Microscopy (AFM)

AFM was performed using the JPK NanoWizard V (RRID:SCR_017787) in PeakForce QNM Tapping mode with qp-BioAC (CB3) cantilevers (NanoAndMore; C = 0.06 N/m; f_0_ = 30 kHz) in DI water. For sample preparation, mica substrates were incubated with 200 μL vesicles at 40 °C for 1 h, then washed with 8 mL DI water. NS5B (100 nM) was applied and incubated 30 min, followed by washing and re-imaging. RNA (100 nM) was added and treated similarly. AFM height data were processed in JPK software and analyzed in Gwyddion. Images were leveled, background-subtracted, FFT-filtered, and masked to extract membrane and NS5B features.

## Acknowledgements

We thank Karla Kirkegaard and Theresia Reindel for critical reading of the manuscript, and the members of the Yang lab for helpful discussions and support. ChatGPT by OpenAI was used for text refinement and for the generation of illustrations. A portion of this work was performed at Macromolecular Structure Group, Nucleus at Sarafan ChEM-H. This work utilized the NanoTemper Prometheus Panta dynamic light scattering (DLS) and nanoDSF system that was purchased with funding from Stanford c-SHARP Program. Structures are visualized using PyMOL(63). This work was supported by the National Institutes of Health (NIH) under grant number R21AI148972.

