## Supplementary information for "Membrane binding promotes oligomerization and functional activation of viral RNA-dependent RNA polymerase"

### Supplementary Figures

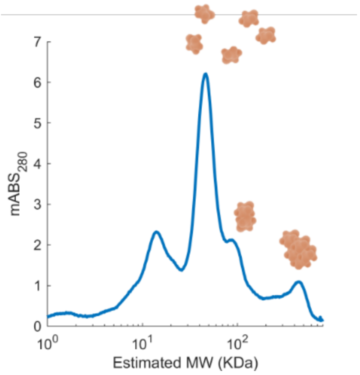


SI Figure 1. NS5B shows as a monomer in size exclusion chromatography. Size-exclusion chromatography (SEC) shows that NS5BΔ21-FlAsH-His is predominantly monomeric at ~1 µM, based on a ~1 ml elution peak. A representative chromatogram from three independent runs is shown; ~200 µg of protein was loaded and compared to a molecular weight standard.

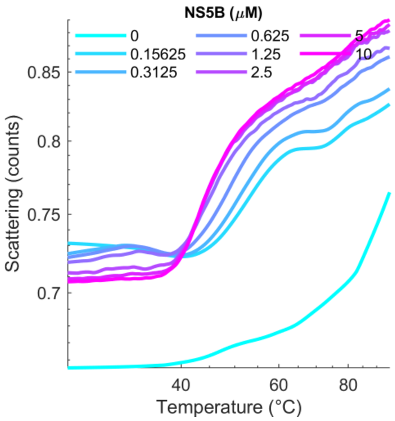


SI Figure 2. nanoDSF shows NS5B melting temperature is above 40º C. Similar results obtained for 2 other independent experiments.

### Supplementary methods

*Size Exclusion Chromatography*

~200 µg of NS5B was run on Superdex 200 Increase 10/300 GL (Cytiva) in buffer (400 mM NaCl, 50 mM NaPi, 5 mM $MgCl_2$, 4 °C) at 0.25-0.3 mL/min. Standards (Supelco) were run for calibration.

*Lower bound estimate for membrane tethered NS5B oligomerization dissociation constant*

Consider that in 1 min observation no recovery was observed for NS5B oligomers. This implies that the probability (P) for lack of dissociation in 60 s is almost certain. Assuming first order reaction for de-oligomerization, we set: P(t = 60 s) > 0.99 > $e^{-k_{off}\times 60\ s}$. Hence $k_{off}$< 1.6× $10^{-4}$ $s^{1}$. For simplicity, assuming 3D diffusion-mediated on-rate of 0.1 $\mu M^{-1}s^{-1}$ we get the dissociation constant to be < 1.6 nM, or by rounding to the nearest upper order of magnitude, <10 nM.

*Estimated protein-membrane absorption in flow channel and in wells*

A single lipid headgroup occupies approximately 0.6 $nm^2$, corresponding to ~6 × $10^{11}$ Ni-NTA headgroups in one leaflet of a 3 mm diameter membrane. Assuming a flow chamber volume of 2.5 µL, the membrane is expected to become saturated with protein at ~400 nM. In comparison, for wells containing 50 μL of solution, saturation should occur at protein concentrations in the tens of nanomolar range.

*TIRF Microscopy*

Imaging done by CellTIRF Olympus system equipped with UPLAPO60XOHR 60× objective, quad filter cube, and Oxxius lasers (405, 488, 561, 648 nm). Imaging used an ORCA-Fusion-BT camera with a W-View Gemini beam splitter. Temperature was controlled with a Tokai-Hit ThermoBox.
